# Non-proliferative adult neurogenesis in neural crest-derived stem cells isolated from human periodontal ligament

**DOI:** 10.1101/325613

**Authors:** Carlos Bueno, Marta Martínez-Morga, Salvador Martínez

**Affiliations:** Instituto de Neurociencias de Alicante (UMH-CSIC), San Juan, Alicante, 03550, Spain; Department of Human Anatomy and Institute of Biomedical Research (IMIB), University of Murcia, Faculty of Medicine, Murcia, 30800, Spain

**Author notes:** Corresponding author: Carlos Bueno, PhD., Instituto de Neurociencias de Alicante. UMH-CSIC, Campus de San Juan, E-03550-Alicante, Spain.

**Keywords:** Neuronal polarity, neural stem cells, adult stem cells, neural crest stem cells, periodontal ligament stem cells, nuclear remodeling, micronuclei

## Abstract

Understanding the sequence of events from undifferentiated stem cells to neuron is not only important for the basic knowledge of stem cell biology, but also for therapeutic applications. The aim of the present study is to evaluate the sequence of biological events occurring during neural differentiation of human periodontal ligament stem cells (hPDLSCs). Here, we show that hPDLSCs-derived neurons display a sequence of morphologic development highly similar to those reported before in primary neuronal cultures derived from rodent brains. We observed that cell proliferation is not present through neurogenesis from hPDLSCs. Futhermore, we may have discovered micronuclei movement and transient cell nuclei lobulation coincident to in vitro neurogenesis. Morphological analysis also reveals that neurogenic niches in the adult mouse brain contains cells with nuclear shapes highly similar to those observed during *in vitro* neurogenesis from hPDLSCs. Our results provide additional evidence that it is possible to differentiate hPDLSCs to neuron and suggest the possibility that the sequence of events from stem cell to neuron does not necessarily requires cell division from stem cell.

## 1. Introduction

Neural stem cells (NSCs) are multipotent populations of undifferentiated cells present both during development and in the adult central nervous system that give rise to new neurons and glia [1]. The presence of neural stem cells in the adult mammalian brain (aNSCs) have been described in two neurogenic niches, the ventricular-subventricular zone (V-SVZ) of the anterolateral ventricle wall and the subgranular zone (SGZ) of the hippocampal dentate gyrus [2–9].

The study of the cell composition of neurogenic niches and the use of methods for detecting proliferating cells, suggest that neurogenesis occurs progressively through sequential phases of proliferation and the neuronal differentiation of aNSCS.

In the V-SVZ, putative aNSCs (type B cells) divide to give rise to intermediate progenitor cells (type C cells), which divide a few times before becoming neuroblasts (type A cells). The neuroblast then migrate into the olfactory bulb and differentiate into distinct types of neurons [2–4]. In the SGZ, putative aNSCs (type 1 cells) divide to give rise to intermediate progenitor cells (type-2 cells) which exhibit limited rounds of proliferation before generating polarized neuroblast (type-3 cells) [5–9]. Neuroblast, as polarized cells, then migrate, guided by the leading process, along SGZ and differentiate into dentate granule neurons [10,11].

However, there are almost no studies that show mitotic chromosomes or mitotic spindle to really confirm that neurogenesis occurs progressively through sequential phases of proliferation and the neuronal differentiation of aNSCS [2–9]. In addition, the selfrenewal and multipotent properties demonstrated by NSC *in vitro* [12] have not been clearly demonstrated *in vivo* [10,13,14].

Ultrastructure and immunocytochemistry studies show that the V-SVZ stem cell niche contains cells with irregular (polymorphic) nuclei [15–17]. Type-B cells have irregular nuclei that frecuently contain invaginations. Type-C cells nuclei contain deep invaginations and Type-A cell nuclei are also occasionally invaginated [2]. Futhermore, recent studies have shown that murine and human V-SVZ contains cells with segmented nuclei connected by an internuclear bridge [18–20]. Although it has been suggested that these are associated with quiescence in aNSCs [20], the functional significance of different nuclear morphologies remains elusive.

Ultrastructure and immunocytochemistry studies also show that the SGZ stem cell niche contains cells with irregular (polymorphic) nuclei [21–28]. Type-2 cells had an irregularly shaped nucleus [7,9]. Importantly, one study found that many cultured hippocampal neurons have irregular nuclei or even consisted of two or more lobes connected by an internuclear bridge [29].

Moreover, how neuroblasts acquire the appropriate cell polarity to initiate their migration remains unclear [30]. The process of neuronal polarization has been studied for decades using dissociated rodent embryonic hippocampal pyramidal neurons and postnatal cerebellar granule neurons in culture [31,32]. During neuronal polarization *in vitro,* the morphological changes in cultured neurons are divided into different stages.

Upon isolation, dissociated pyramidal neurons retract their processes, so that their development *in vitro* begins as rounded spheres that spread lamellipodia (stage 1). These spheres appear symmetrical, extending and retracting several immature neurites of a similar length (stage 2). Elongation of a single process, that which presumably becomes the axon, breaks this symmetry (stage 3). The next step involves the remaining short neurites morphologically developing into dendrites (stage 4) and the functional polarization of axon and dendrites (stage 5), including dendritic spine and synapse formation [33]. Dissociated granule neurons also present a lamellipodia after attaching to the substratum (stage 1). These spheres extend a unipolar process at a single site on the plasma membrane (stage 2) followed by extension of a second process from the opposite side of the cell body, resulting in a bipolar morphology (stage 3). One of the two axon elongates futher and start branching (stage 4), and shorter dendritic processes develop around the cell body (stage 5) [34].

Understanding the sequence of events from aNSCs to neuron is not only important for the basic knowledge of NSCs biology, but also for therapeutic applications [35]. The major barrier to studying human aNSCs is the inaccessibility of living tissue, therefore an enormous effort has been made in this study to derive neurons from human stem cells [36]. *In vitro* models of adult neurogenesis mainly utilize fetal, postnatal and adult NSCs [37].

Neural crest stem cells (NCSCs) are a migratory cell population that generate numerous cell lineages during development, including neurons and glia [38,39]. NCSCs are present not only in the embryonic neural crest, but also in various neural crest-derived tissues in the fetal and even adult organs [40]. The periodontal ligament (PDL) is a connective tissue surrounding the tooth root that contains a source of human NCSCs which can be accessed with minimal technical requirements and little inconvenience to the donor [41]. Isolation and characterization of multipotent stem cells from the human PDL have been previously described [42,43].

In previous publication, we showed that several stem cell and neural crest cell markers are expressed in human adult periodontal ligament (hPDL) tissue and hPDL-derived cells. [44]. *In vitro,* hPDL-derived cells differentiate into neural-like cells based on cellular morphology and neural marker expression. *In vivo,* hPDL-derived cells survive, migrate and expressed neural markers after being grafted to the adult mouse brain. Futhermore, some hPDL-derived cells graft into stem cell niches such as V-SVZ of the anterolateral ventricle wall and the SGZ of the dentate gyrus in the hippocampus. It is important to mention that the hPDLSCs located in the stem cell niches show neural stem morphology. Moreover, hPDLSCs expressed ion channel receptors [45] and displayed inward currents conducted through voltage-gated sodium (Na+) channels and spontaneous electrical activities after neurogenic differentiation [46,47]. Therefore, the neural crest origin and neural differentiation potential both *in vitro* and *in vivo*, make human periodontal ligament stem cells (hPDLSCs) interesting for investigating stem cell to neuron differentiation mechanisms.

Here, we show that hPDLSCs-derived neurons display a sequence of morphologic development highly similar to those reported before in primary neuronal cultures derived from rodent brains during neurogenesis, providing additional evidence that it is possible to differentiate hPDLSCs to neuron.

We observed that cell proliferation is not present through neurogenesis from hPDLSCs. In fact, the cell shape of hPDLSCs is reset and start their neuronal development as round spheres. Futhermore we may have discovered a transient cell nuclei lobulation coincident to *in vitro* neurogenesis, without being related to cell proliferation. We observed that small DNA containing structures may move within the cell to specific directions and temporarily form lobed nuclei.

Morphological analysis also reveals that the V-SVZ of the anterolateral ventricle wall and the SGZ of the hippocampal dentate gyrus in the adult mouse brain contains cells with nuclear shapes highly similar to those observed during *in vitro* neurogenesis from hPDLSCs. We suggest the possibility that neuronal differentiation from aNSCs may also occur during *in vivo* adult mammalian neurogenesis without being related to cell proliferation.

## 2. Materials and methods

### 2.1. Ethical conduct of research

Methods were carried out in accordance with the relevant guidelines and regulations. The experimental protocols were approved by the Institutional Review Board of the Miguel Hernández University of Elche (No. UMH.IN.SM.03.16) and the signed informed consent was obtained from all patients before the study. The authors declare that all experiments on human subjects were conducted in accordance with the Declaration of Helsinki. All protocols and care of the mice were carried out according to the guidelines of the European Communities Council Directive of 24 November 1986 (86/609/EEC). The authors further attest that all efforts were made to minimize the number of animals used and their suffering.

### 2.2. Cell Culture

Human premolars were extracted and collected from healthy adult donors undergoing orthodontic therapy in Murcia dental hospital (Spain). hPDL was scraped from the middle third region of the root surface. After washing the extracted PDL with Ca and Mg-free Hank’s balance salt solution (HBSS; Gibco), hPDL was digested with 3 mg/ml type I collagenase (Worthington Biochemical Corporation) and 4 mg/ml dispase II (Gibco) in alpha modification minimum essential medium eagle (α-MEM) (α-MEM; Sigma-Aldrich) for 1 h at 37°C. The reaction was stopped by the addition of α-MEM. The dissociated tissue was passed through a 70-μm cell strainer (BD Falcon). Cells were centrifuged, and the pellet was resuspended in in serum-containing media (designated as the basal media), composed of α-MEM supplemented with 15% calf serum (Sigma), 100 units/ml penicillin-streptomycin (Sigma) and 2 mM l-glutamine (Sigma). The cell suspension was plated into six-well multiwell plates (BD Falcon) and incubated at 37°C in 5% CO2. To induce neural differentiation, cells were cultured in serum-free media (designated as the neural induction media), consisting in Dulbecco’s modified Eagle’s medium/F12 (DMEM/F12, Gibco) supplemented with bFGF (20 ng/ml, R&D Systems), EGF (20 ng/ml, R&D Systems), glucose (0.8 mg/ml, Sigma), N2-supplement (Gibco), 2 mM l-glutamine (Sigma), and 100 units/ml penicillin-streptomycin (Sigma). Neural induction media were changed every 3-4 days until the end of the experiment (2 weeks).

### 2.3. Immunocytochemistry

Cells were plated onto coated plastic or glass coverslips, and maintained in basal media or neural induction media. Cells were rinsed with PBS and fixed in freshly prepared 4% paraformaldehyde (PFA; Sigma). Fixed cells were blocked for 1 h in PBS containing 10% normal horse serum (Gibco) and 0.25% Triton X-100 (Sigma) and incubated overnight at 4°C with antibodies against: β-III-tubulin (TUJ1; 1:500, Covance), Tau (GTX49353; 1/300, GeneTex), MAP2 (840601; 1/300, Biolegend), Connexin-43 (3512; 1/300, Cell Signalling), Synaptophysin (18-0130; 1/300, Zymed), Synapsin1 (NB300-104; 1/300, Novus), Fibrillarin (ab5821; 1/300, Abcam) and Laminin A/C (GTX101127; 1/300, GeneTex) in PBS containing 1% normal horse serum and 0.25% Triton X-100. On the next day, cells were rinsed and incubated with the corresponding secondary antibodies: Alexa Fluor^®^ 488 (anti-mouse or anti-rabbit; 1:500, Molecular Probes), Alexa Fluor^®^ 594 (anti-mouse or anti-rabbit; 1:500, Molecular Probes), biotinylated anti-rabbit (BA1000, 1:250; Vector Laboratories), biotinylated anti-chicken (BA9010, 1:250, Vector Laboratories, CY3-streptavidin (1:500, GE Healthcare). Cell nuclei were counterstained with DAPI (0.2 mg/ml in PBS, Molecular Probes). Alexa Fluor 488^®^ phalloidin was used to selectively stains F-actin (Molecular Probes).

### 2.4. Western Blotting

hPDL-derived cells were harvested using trypsin/EDTA (Gibco), washed twice with PBS, resuspended in RIPA lysis buffer (Millipore) for 30 min at 4°C in the presence of protease inhibitors (Pierce™. protease inhibitor Mini Tables, Pierce Biotechnology Inc) and PMSF 1M (Abcam). Protein concentration was determined using the bradford protein assay (Sigma-Aldrich). Proteins were separated in 8% SDS-polyacryamide gel (PAGE-SDS) and transferred to a nitrocellulose membrane (Whatman). PageRuler™ Prestained Protein Ladder (Thermo Scientific) has been used as size standards in protein electrophoresis (SDS-PAGE) and western-blotting. After transfer, nitrocellulose membranes were stained with Ponceau S solution (Sigma-Aldrich) to visualize protein bands. Blots were then incubated over-night at 4°C with rabbit antibody against β-III-tubulin (TUJ1; 1:1000, Covance). Secondary antibody was used at 1:7000 for peroxidase anti-mouse Ab (PI-2000, Vector Laboratories). Immunoreactivity was detected using the enhanced chemiluminescence (ECL) Western blot detection system (Amersham Biosciences Europe) and Luminata™ Forte (Millipore corporation) using ImageQuant *LAS 500* Gel Documentation System (GE Healthcare). The molecular weight of β-III-tubulin is approximately 55 kDa.

### 2.5. Immunohistochemistry

Experiments were carried out according to the guidelines of the European Community (Directive 86/609/ECC) and in accordance with the Society for Neuroscience recommendations. Animals used in this study were 12-week-old immune-suppressed mouse (Hsd:Athymic Nude-Foxn1 nu/nu; Harlan Laboratories Models, S.L), housed in a temperature and humidity controlled room, under a 12h light/dark cycles, with *ad libitum* access to food and water. The animals were anesthetized and intracardially perfused with freshly prepared, buffered 4% PFA (in 0.1M PB, pH 7.4). Brains were removed, postfixed for 12 hr in the same fixative at 4°C and dehydrated in 30% sucrose solution at 4°C until sunk. 30μm thick coronal sections were collected using a freezing microtome. Serial sections were used for DAPI staining. Free-floating sections were incubated and mounted onto Superfrost Plus glass slides (Thermo Scientific). The slides were dried O/N and coverslipped with mowiol-NPG (Calbiochem).

### 2.6. Images and Data Analyses

Analyses and photography of visible and fluorescent stained samples were carried out in an inverted Leica DM IRB microscope equipped with a digital camera Leica DFC350FX (Nussloch) or in confocal laser scanning microscope Leica TCS-SP8. Digitized images were analyzed using LASX Leica confocal software. Z-stacks of confocal fluorescent images were also analyzed to calculate the nuclear volume by using ImageJ software.

### 2.7. Scanning Electron Microscopy

Cells were plated onto coated glass coverslips and maintained in basal media or neural induction media. Cells were treated with fixative for 20 minutes. Coverslips were postfixed in 1% osmium tetroxide for 1 hour and dehydrated in graded ethanol washes. The coverslips were allowed to dry at a conventional critical point and were then coated with gold-palladium sputter coated. Coverslips were view on a Jeol 6100 scanning electron microscope.

### 2.8. Transmission Electron Microscopy

hPDL-derived cells were harvested using trypsin/EDTA (Gibco) and were treated with fixative for 60 minutes. Cells were postfixed in osmium tetroxide solution, dehydrated embedded in resin. Ultrathin sections (70–90 nm) were cut, stained with lead citrate, and examined under Jeol 1011 and Philips Tecnai 12 transmission electron microscopes.

## 3. Results

As noted in the introduction, the aim of this work was to evaluate the sequence of biological events occurring during the neural differentiation of hPDLSCs. Morphological characteristics of the hPDLSCs, including cell shape, cell surface features, cytoskeleton, and nuclear morphology were examined in cells under proliferation and neural differentiation conditions.

### 3.1 hPDLSCs cultured in basal media

Under proliferation conditions, hPDLSCs displayed a fibroblast-like morphology with low-density microvilli on the cell surface (Figure 1A) and actin microfilaments and β-III tubulin microtubules oriented parallel to the longitudinal axis of the cell (Figure 1B). The cytoskeletal protein class III beta-tubulin isotype is widely regarded as a neuronal marker in developmental neurobiology and stem cell research [48]. Dental and oral-derived stem cells displayed spontaneous expression of neural marker β-III tubulin, even without having been subjected to neural induction [49]. Western blot analysis verified the expression of β-III tubulin in hPDLSCs (Figure 1C). During mitosis, β-III tubulin is present in the mitotic spindle and it is detectable in all phases of mitosis (Figure 1D). The cytoskeletal protein class III beta-tubulin isotype is a component of the mitotic spindle in multiple cell types [50]. During interphase, undifferentiated hPDLSCs displayed a flattened, ellipsoidal nucleus, often located in the center of the cell and with a nuclear volume around 925’356 ±52’6184 μm^3^ (Figure 1E).

**Figure 1.**
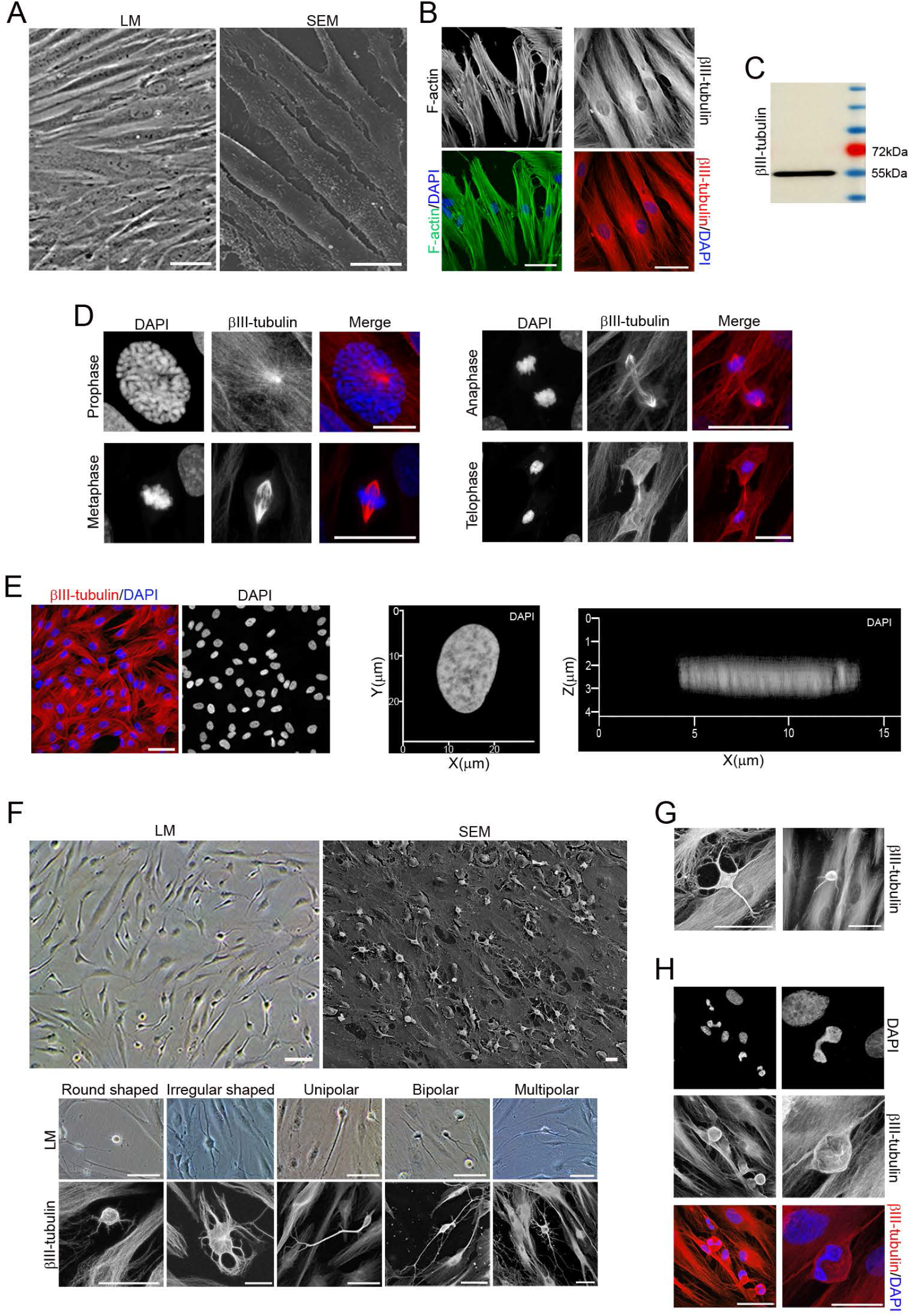
Morphological changes in hPDLSCs cultures during neural induction. Undifferentiated hPDLSCs presented a fibroblast-like morphology with low-density microvilli on their surface (**A**) and actin microfilaments and β-III tubulin microtubules oriented parallel to the longitudinal axis of the cell (**B**). (**C**) Western blot analysis verified the expression of β-III tubulin. Protein size markers (in kilodaltons) are indicated on the side of the panel. (**D**) During mitosis, β-III tubulin is present in the mitotic spindle and it is detectable in all phases of mitosis. (**E**) Undifferentiated hPDLSCs displayed a flattened, ellipsoidal nucleus often located in the center of the cell. (**F**) After 14 days of neural differentiation conditions, hPDLSCs with different morphologies were observed. (**G**) In addition, hPDLSCs of various size were also observed. (**H**) Microscopic analysis also revealed that some hPDLSCs have different nuclear size and shapes, including lobed nuclei connected by an internuclear bridge. Scale bar: 25 μm. LM, light microscopy; SEM, scanning electron microscopy.

### 3.2. hPDLSCs cultured in neural induction media

After 14 days of neural differentiation conditions, the hPDLSCs displayed different morphologies, including round cells with small phase-bright cell bodies and short processes; highly irregulary-shaped cells; and, also, unipolar, bipolar and multipolar-shaped cells with small phase-bright cell bodies and multiple branched processes (Figure 1F). In addition, cells of different size were also observed (Figure 1G). Futhermore, microscopic analysis revealed that some hPDLSCs have different nuclear shapes, including lobed nuclei connected by an internuclear bridge (Figure 1H). The results may indicate that the cell culture simultaneously contains hPDLSCs at different stages of neurogenesis and neuronal polatization. We acknowledge that the definitive sequence of *in vitro* neurogenesis and neuronal polarization from hPDLSCs will be provided only by time-lapse microscopy of a single cell, but in our experimental conditions, several pieces of data suggest how these steps may occur.

### 3.3. In vitro neurogenesis from hPDLSCs

After neural induction, hPDLSCs undergo a dramatic change in shape and size, first adopting highly irregular forms and then gradually contracting into round cells with small phase-bright cell bodies (Figure 2A). Cytoskeletal remodeling is observed during the morphological changes that occurred when the hPDLSCs round up to a near-spherical shape. Actin microfilament not longer surround the nucleus and became cortical. Unlike actin, β-III tubulin seems to accumulate around the nucleus (Figure 2B). Actin microfilament and β-III tubulin microtubule network are almost lost in the rounded cells (Figure 2C). Scanning electron micrographs show that hPDLSCs also experience dramatic changes in cell surface features. Under proliferation conditions, hPDLSCs remain very flat, presenting low-density microvilli on their surface (Figure 1A), but there is a marked increase in the number of microvilli as the cells round up to near-spherical shape (Figure 2D). The surface of the round cells is almost devoid of microvilli (Figure 2E). Cytokinesis and mitotic spindle were not observed during the described of *in vitro* neurogenesis processes (Figure 1F-H,2).

**Figure 2.**
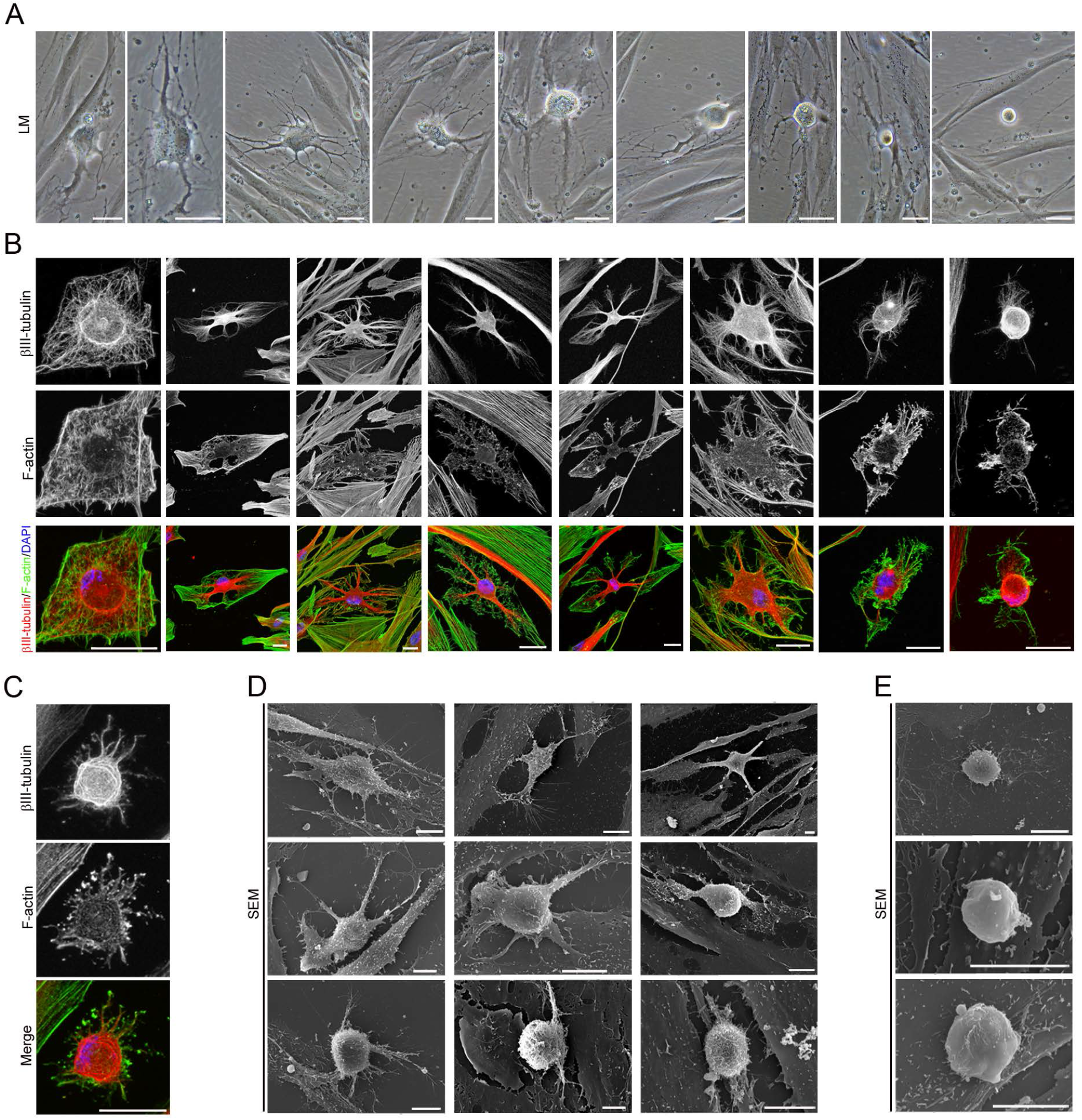
*In vitro* neurogenesis from hPDLSCs. (**A**) After neural induction, hPDLSCs undergo a shape and size change, adopting highly irregular forms first and then gradually contracting into round cells. (**B**) Cytoskeletal remodeling is observed during these morphological changes. Actin microfilament not longer surround the nucleus and become cortical. Unlike actin, β-III tubulin seems to accumulate around the nucleus. (**C**) the cytoskeletal network is almost lost in round cells. (**D**) Scanning electron micrographs show that there is a marked increase in the density of microvilli as the cells round up to near-spherical shape. (**E**) The surface of round cells is almost devoid of microvilli. The scale bars are 25 μm in the light microscope images, and 10 μm in the scanning electron micrographs. LM, light microscopy; SEM, scanning electron microscopy.

### 3.4. Neuronal polarization of hPDLSCs-derived neurons

Morphological analysis revealed that hPDLSCs-derived neurons display a sequence of morphologic development highly similar to those reported before in dissociated-cell cultures prepared from rodent brain (Figure 3–5). hPDLSCs-derived neurons also start their development as rounded spheres that initiated neurite outgrowth at a single site on the plasma membrane, first becoming unipolar, stages 1-2 (Figure 3A). We did not observe the development of lamellipodia around the circumference of the cell body. These unipolar cells, later transformed into cells containing several short neurites, developed around the cell body, stage 3 (Figure 3B). An analysis of the cytoskeletal organization during spherical stages of hPDLSCs-derived neurons showed that the β-III tubulin microtubules and actin microfilament network is reorganized. Cytoskeletal protein β-III tubulin was densely accumulated under the cell membrane of the hPDLSCs-derived neurons cell bodies and in cell neurites (Figure 3A,B) while actin microfilaments were mainly found in cell neurites (Figure 3C). We observed that hPDLSCs-derived neurons produce neurites that showed growth cone formations at their tips (Figure 3C-E). The central domain of the growth cone contains β-III tubulin microtubules and the peripheral domain is composed of radial F-actin bundles (Figure 3D), similar to the typical spatial organization described in neurons [51,52]. Scanning electron micrographs also showed that the growth cone of hPDLSCs-derived neurons contained filopodia and vesicles on the cell surface (Figure 3E). These finding are consistent with a previous study reporting that membrane addition and extension in growth cones is mediated by diverse mechanism, including exocytosis of vesicular components [53]. Microtubule-associated proteins Tau and MAP2 were also found in hPDLSCs-derived neurons (Figure 3F). At later stages of differentiation, the hPDLSCs-derived neurons gradually adopted a complex morphology by forming several processes, stage 4 (Figure 3G) that grew and arborized, adquiring dendritic-like and axonal-like identities, giving rise to a variety of neuron-like morphologies (Figure 3H). The next step, stage 5, in neuronal polarization from rodent neurons in culture is the functional polarization of axon and dendrites, including dendritic spine formation and axon branch formation. Dendritic spines are micron-sized dendrite membrane protrusions [54]. Depending on the relative sizes of the spine head and neck, they can be subdivided into different categories, including filopodium, mushroom, thin, stubby, and branched spines [55]. Dendritic spines are actin-rich compartments that protrude from the microtubule-rich dendritic shafts of principal neurons [56]. Based on morphology, complexity, and function, axon branching is grouped into different categories, including arborization, bifurcation, and collateral formation [57].

**Figure 3.**
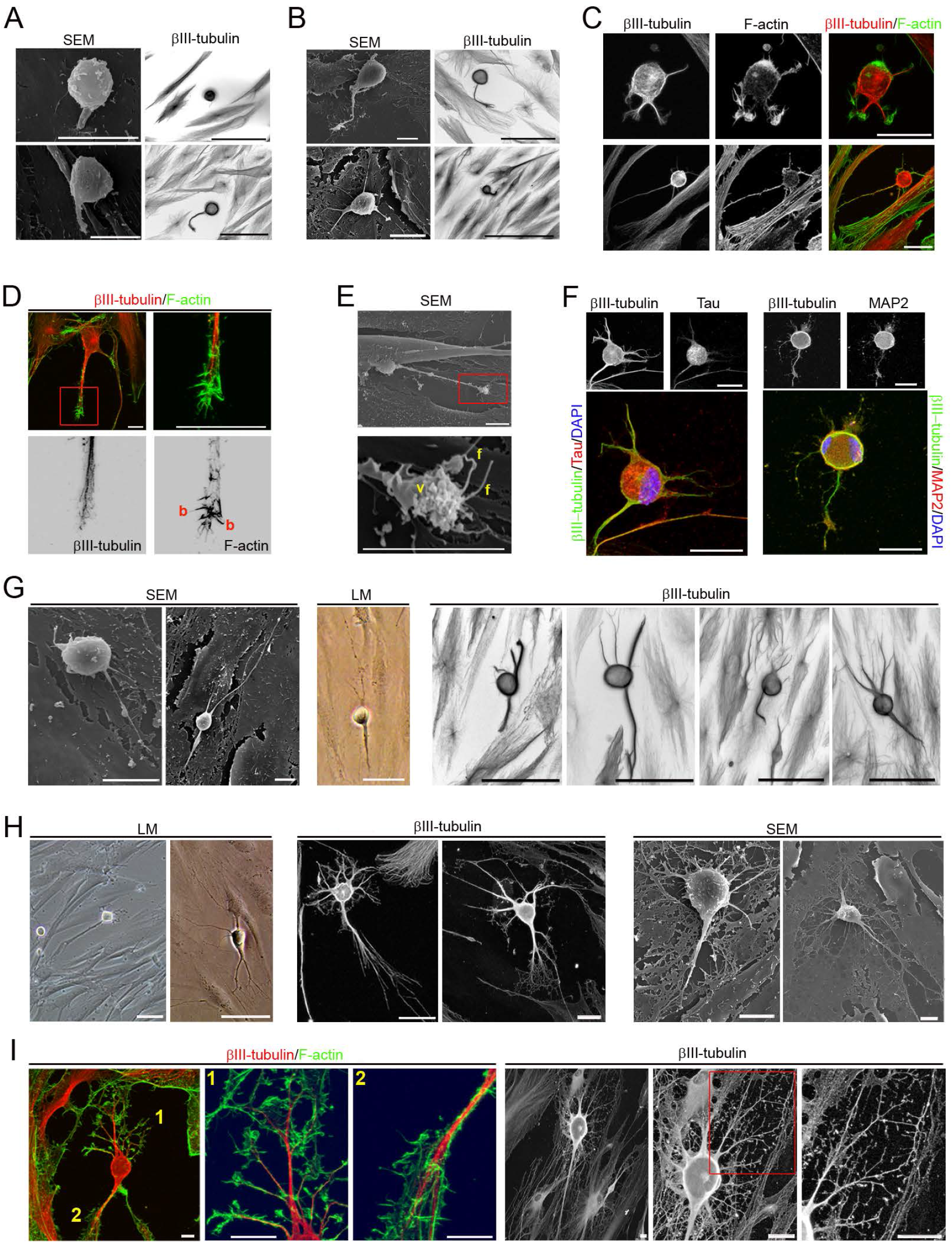
Neuronal polarization of hPDLSCs-derived neurons. (**A**) hPDLSCs-derived neurons start their development as rounded spheres that initiate neurite outgrowth at a single site on the plasma membrane. (**B**) These later transform into cells containing several short neurites developed around the cell body. (**C**) The cytoskeletal network is reorganizated. β-III tubulin accumulates densely under the cellular membrane of the cell body and in cell neurites while actin microfilaments are mainly found in cell neurites. (**D**) The peripheral domain in the growth cone of hPDLSCs-derived neurons is composed of radial F-actin bundles and the central domain contains β-III tubulin microtubules. (**E**) Micrographs showing that the growth cone also contains filopodia and vesicles on the cell surface. (**F**) Microtubule-associated proteins Tau and MAP2 were also found in hPDLSCs-derived neurons. At later stages of development, hPDLSCs-derived neurons gradually adopt a complex morphology (**G**) giving rise to a variety of neuron-like forms (**H**). (**I**) Cytoskeletal protein β-III tubulin and F-actin staining shown that hPDLSCs-derived neurons develop distinct axon-like and dendrite-like processes (numbers locate the areas shown in higher power). The scale bars are 25 μm in the light microscope images, and 10 μm in the scanning electron micrographs. b, actin bundles; f, filopodia; LM, light microscopy; SEM, scanning electron microscopy; v, vesicles.

Our morphological analysis revealed that hPDLSCs-derived neurons developed well-differentiated axonal-like and dendritic-like domains. These types of processes differ from each other in morphology (Figure 3I–4D). Cytoskeletal protein β-III tubulin and F-actin staining showed that the hPDLSCs-derived neurons comprised multiple branched dendrite-like processes with dendritic spines-like structures (Figure 3I). Scanning electron micrographs showed that the hPDLSCs-derived neurons also contained multiple branched dendrite-like processes with variously shaped spine-like protusions, highly similar to filopodium, mushroom, thin, stubby, and branched dendritic spines shapes (Figure 4A). Futhermore, hPDLSCs-derived neurons also displayed different types of axonal branch-like structures, including bifurcation (Figure 4B), arborization (Figure 4C), and collateral formation (Figure 4D).

**Figure 4.**
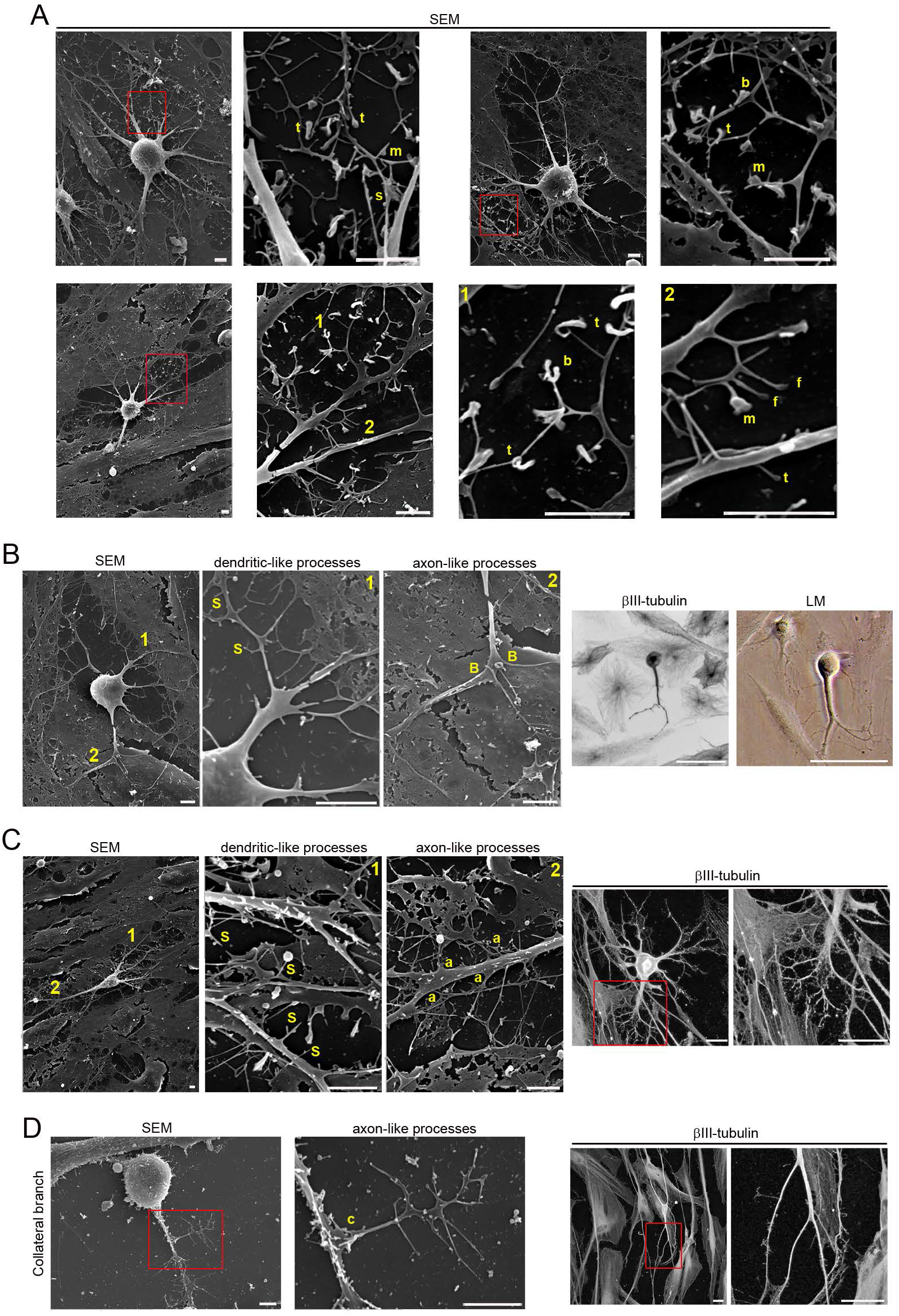
hPDLSCs-derived neurons have developed well-differentiated axonal-like and dendritic-like domains. (**A**) Scanning electron micrographs show that hPDLSCs-derived neurons are composed of multiple branched processes with different spine-like protusions highly similar to filopodium, mushroom, thin, stubby, and branched dendritic spines shapes. hPDLSCs-derived neurons also display different types of axonal branch-like structures, including bifurcation (**B**), terminal arborization (**C**), and collateral formation (**D**) (inserts and numbers locate the areas showed in higher power). The scale bars are 25 μm in light microscope images and 5 μm in the scanning electron micrographs. a, arborization; B, bifurcation; b, branched; c, collateral formation; f, filopodium; LM, light microscopy; m, mushroom; S, spine-like protusions; s, stubby; SEM, scanning electron microscopy; t, thin.

The last step in neuronal polarization from rodent neurons in culture is synapse formation. The most frequent types of synaptic communication include axodendritic, axosomatic, axoaxonic and dendrodendritic synapses. Morphological analysis revealed that the hPDLSCs-derived neurons connected to one another (Figure 5A) through different types of synapse-like interactions, including dendrodendritic-like, axoaxonic-like and axodendritic-like synapses (Figure 5B). Synapse-associated proteins Cx43, Synaptophysin and Synapsin1 were found accumulated in the cell surface of neurites (Figure 5C).

**Figure 5.**
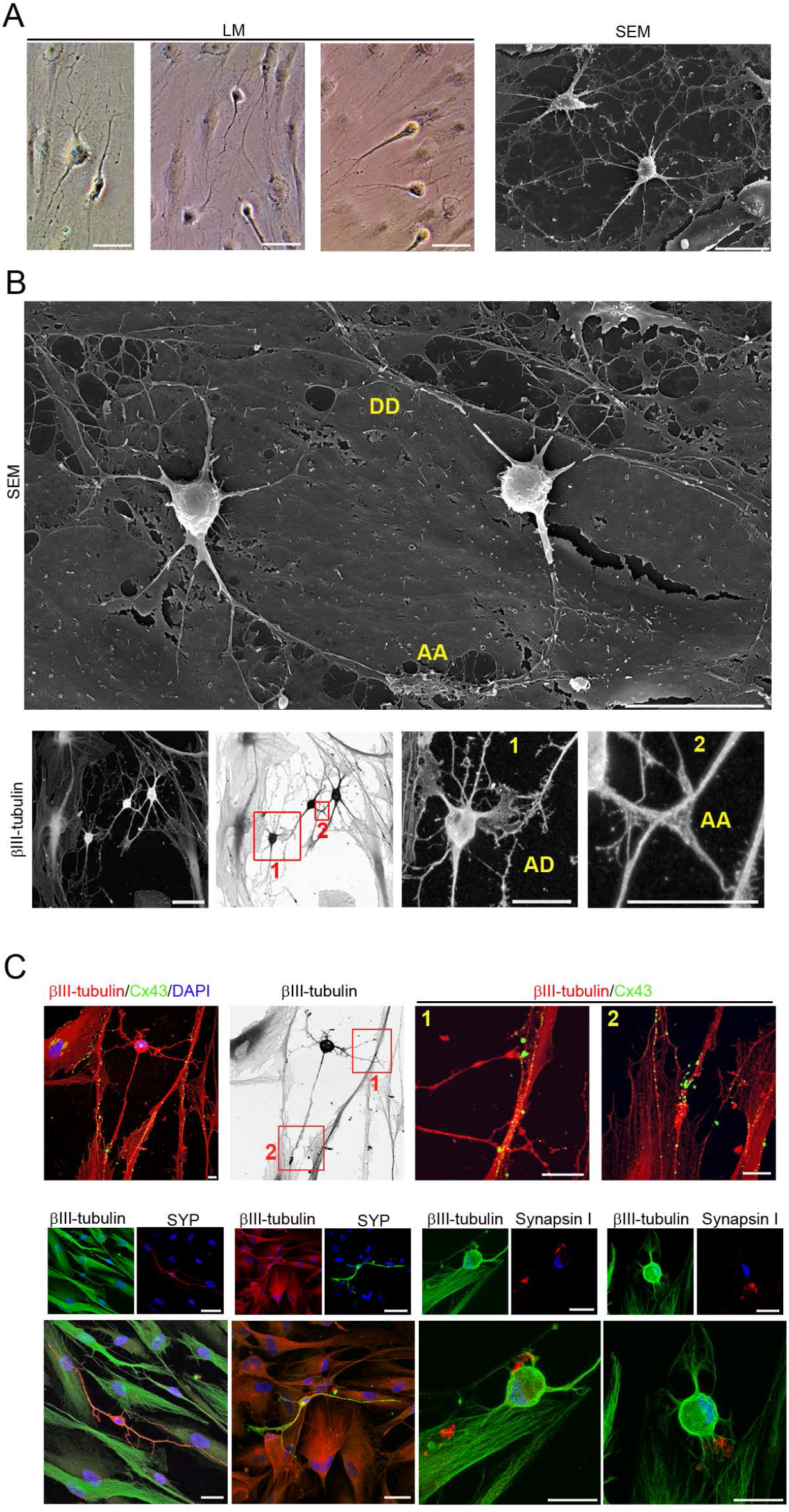
hPDLSCs-derived neurons are connected by synapse-like interactions. hPDLSCs-derived neurons connect to one another (**A**) through different types of synapses-like interactions, including dendrodendritic-like, axoaxonic-like and axodendritic-like synapses (**B**). (**C**) Synapse-associated proteins Cx43, Synaptophysin and Synapsin1 are found in the cell membrane of hPDLSCs-derived neurons at the neurite contact areas. Scale bar: 25 μm. AA, axoaxonic-like synapse; AD, axodendritic-like synapse; DD, dendrodendritic-like synapse; LM, light microscopy.

### 3.5. Nuclear remodeling

Nuclear morphology was examined in hPDLSCs under proliferation and neural differentiation conditions. The dynamic localization of the nucleoli was analyzed by immunostaining for fibrillarin, the main component of the active transcription centers [58] and the dynamic localization of the nuclear lamina was analyzed by immunostaining for laminin A/C, a nuclear lamina component [59].

First, we analyzed the nuclear morphology in proliferative hPDLSCs. As noted above, during interphase, hPDLSCs displayed a flattened, ellipsoidal nucleus, often located in the center of the cell, and with a nuclear volume around 925.356 ±52.6184μm3 (Figure 1E). The nuclei of hPDLSCs contained two or more nucleoli and the inside surface of the nuclear envelope is lined with the nuclear lamina (Figure S1A). Previous studies have shown that the nuclear lamina and nucleolus are reversibly disassembled during mitosis [60,61]. Microscopic analysis of hPDLSCs revealed that the dynamic localization of fibrillarin and laminin A/C proteins during mitosis are similar to those observed in previous studies (Figure S1B). Futhermore, β-III tubulin is present in the mitotic spindle and it is detectable in all phases of mitosis and mitotic chromosomes and cytokinesis were also observed (Figure 1D,S1B).

Morphological analysis also revealed that nuclear remodeling occurred during *in vitro* neurogenesis from hPDLSCs (Figure 6,7). We acknowledge that the definitive sequence of nuclear remodeling when hPDLSCs round up to near-spherical shape will only be provided by time-lapse microscopy, but our accumulated data suggests how these steps may occur.

**Figure 6.**
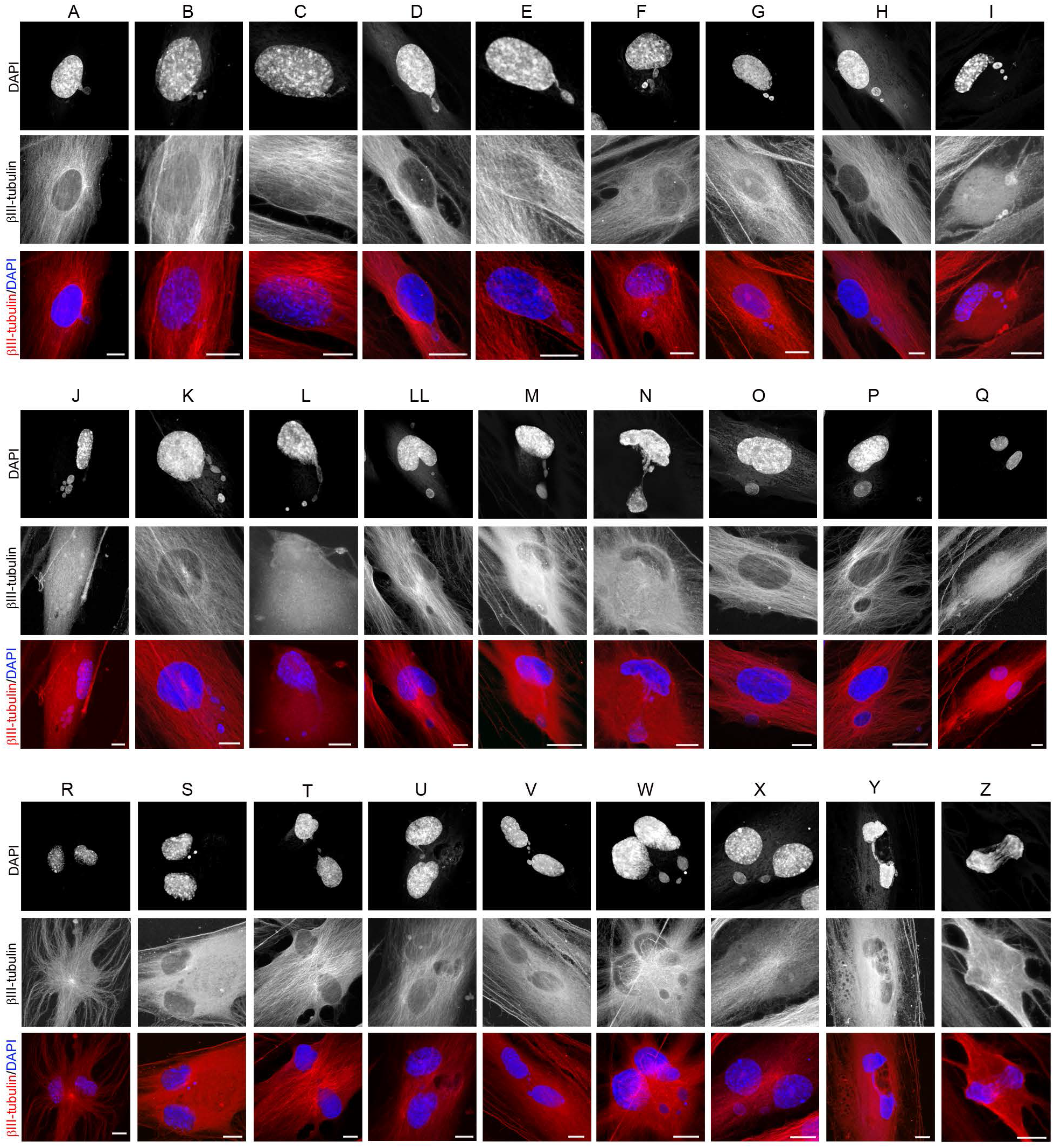
Nuclear shape remodeling occurs during neurogenesis from hPDLSCs. (**A-N**) Small DNA containing structures (micronuclei) arise from the main nuclei (nuclear buds) and start to move towards specific positions within the cell and temporarily form lobed nuclei (**O-R**). Later, these lobed nuclei connected to one another through small structures containing DNA (**S-X**) forming nucleoplasmic bridges (**Y,Z**). Scale bar: 10 μm.

**Figure 7.**
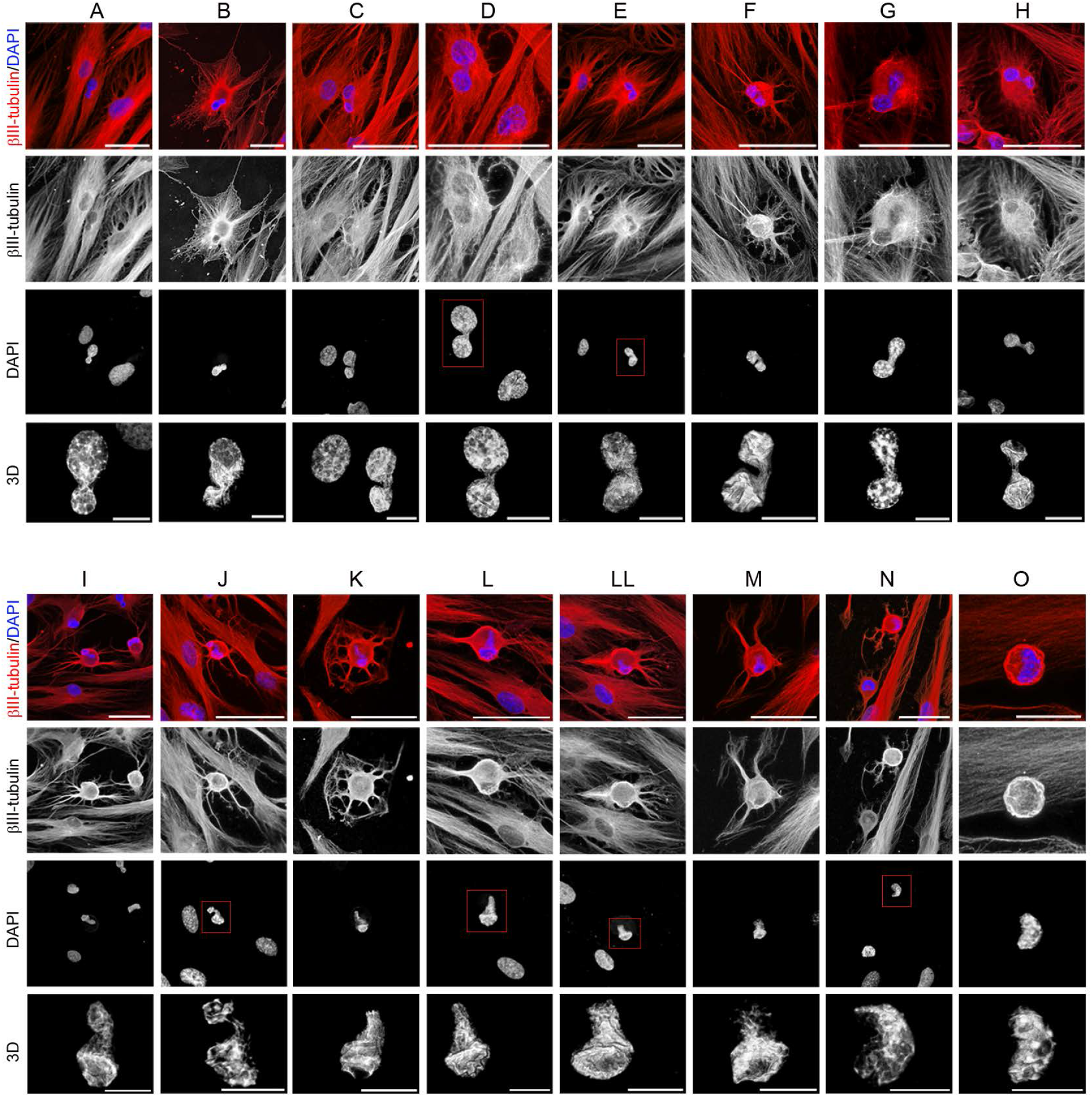
Nuclear shape remodeling occurs during neurogenesis from hPDLSCs. (**A-J**) lobed nuclei connected by nucleoplasmic bridges move towards specific positions within the cell and finally, there is restoration of irregular, but non-lobed, nucleus with an eccentric position within hPDLSCs-derived neurons (**K-O**). The scale bars in β-III tubulin and DAPI images are 50 μm and 10 μm for confocal 3D images of nuclei.

Small DNA containing structures (micronuclei) arise from the main nuclei (nuclear buds) and start to move towards specific positions within the cell (Figure 6A-N) and temporarily form lobed nuclei (Figure 6O-R). Later, these lobed nuclei connected to one another through small DNA containing structures (Figure 6S-X) forming nucleoplasmic bridges (Figure 6Y–7J). Finally, there is restoration of irregular, but non-lobed, nucleus with an eccentric position within hPDLSCs-derived neurons (Figure 7K-O). These small DNA containing structures displayed a spherical or ovoid shape (Figure S2A), and it seems that some of them are connected to the main body of the nucleus by thin strands of nuclear material (Figure S2B). Fibrillarin and laminin A/C proteins were detected in these small DNA containing structures (Figure S2C).

We also observed that the nuclear lamina and nucleolus are not disassembled during *in vitro* neurogenesis from hPDLSCs (Figure S3A). In addition, cytokinesis was not observed during the described of *in vitro* neurogenesis processes from hPDLSCs (Figure S3B). Futhermore, mitotic chromosomes and mitotic spindle were not observed during the described of *in vitro* neurogenesis processes or neuronal polarization from hPDLSCs (Figure 6,7,S2,S3).

In addition, ultrastructural morphological characteristics of hPDLSCs were examined under neural differentiations conditions. Transmission electron microscopy (TEM) is considered the gold standard to confirm apoptosis (62). Apoptotic cell contains certain ultrastructural morphological characteristics, including electron-dense nucleus, disorganized cytoplasmic organelles, large clear vacuoles, nuclear fragmentation and apoptotic bodies. TEM analysis revealed that hPDLSCs under neural differentiations conditions do not meet the criteria described above (Figure S3C). Therefore, hPDLSCs with abnormal nuclei do not represent apoptotic cells.

During neuronal polarization, no lobed nuclei were observed as hPDLSCs-derived neurons gradually acquired a more mature neuronal-like morphology (Figure S4A). We also found that as the cells round up to a near-spherical shape the nuclear volume of the hPDLSCs decreases to an approximate volume of 279.589±38.8905 μm3 (Figure S4B).

Interestingly, the morphological analysis revealed that the adult rodent V-SVZ of the anterolateral ventricle wall (Figure 8A) and the SGZ of the hippocampal dentate gyrus (Figure 8B), where adult neurogenesis has been clearly demonstrated, contained cells with nuclear shapes highly similar to those observed during *in vitro* neurogenesis from hPDLSCs. Although it has been suggested that lobed nuclei connected by an internuclear bridge are associated with quiescence in aNSCs [20], we observed that this kind of nuclei may be associated to nuclear movement within the cell during initial phases of neurogenesis, without being related to cell proliferation.

**Figure 8.**
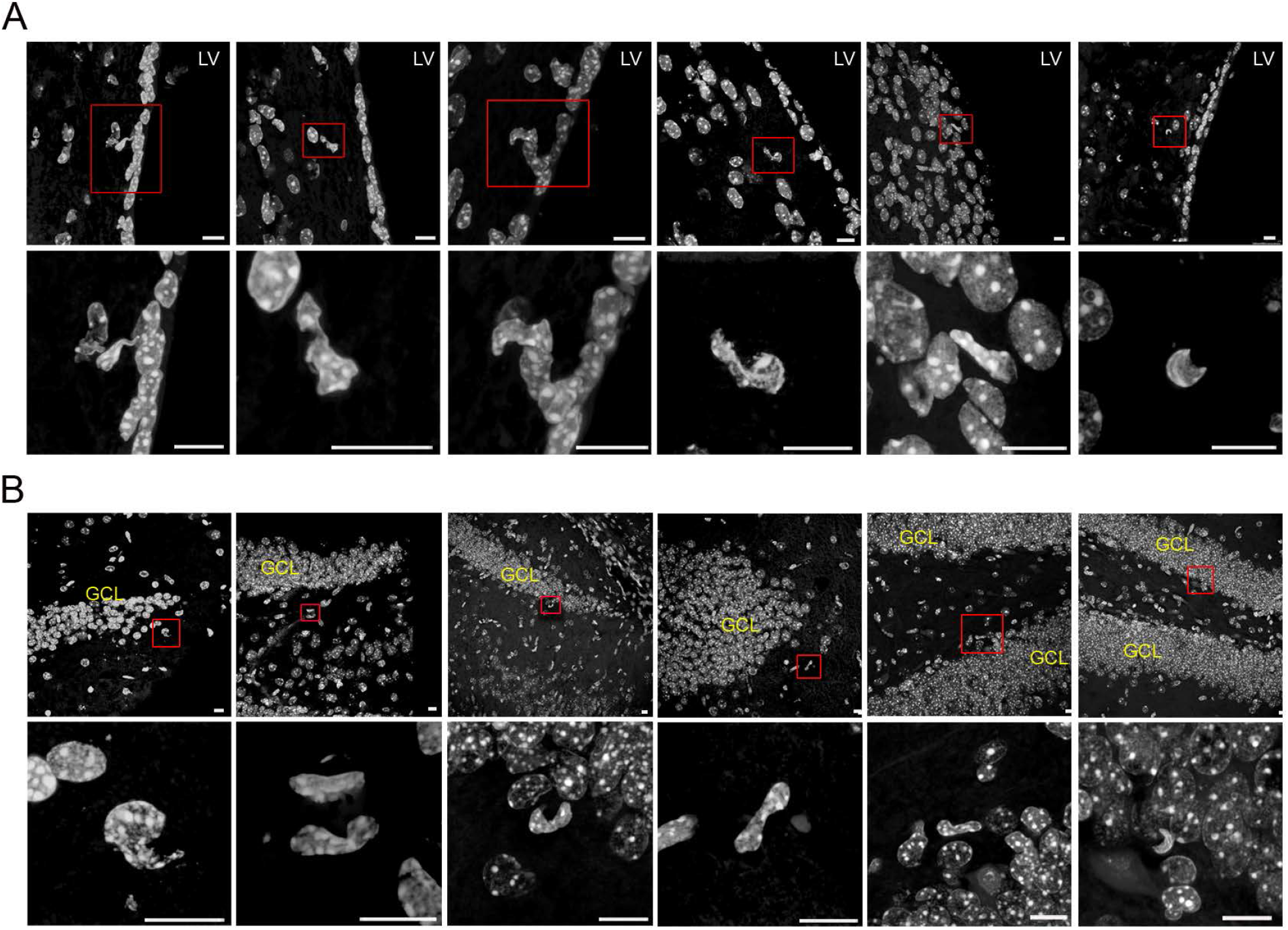
Neurogenic niches in the adult mammalian brain also contains cells with irregular nuclei. Morphological analysis reveals that the adult rodent V-SVZ of the anterolateral ventricle wall (**A**), as well as the SGZ of the hippocampal dentate gyrus (**B**), contain cells with nuclear shapes highly similar to those observed in during in vitro neurogenesis from hPDLSCs. Scale bar: 10 μm. GLC, granule cell layer; LV, lateral ventricle.

## 4. Discussion

The process of neuronal polarization has been studied for decades using dissociated rodent embryonic hippocampal pyramidal neurons and postnatal cerebellar granule neurons in culture [31,32], but less is known about the process of neuronal polarization in human cells [37,63].

In this study we show that hPDLSCs-derived neurons display a sequence of morphologic development highly similar to those reported before in primary neuronal cultures derived from rodent brains. Although future research is required to optimize the diversity of *in vitro* neural induction protocols that have been designed for oral and dental stem cells [64–66], our results provide additional evidence that it is possible to differentiate hPDLSCs to neuron, as suggested by their neural-crest origin, stem cell characteristics and neural differentiation potential both *in vitro* and *in vivo* [64–66, 39–47]. Therefore, we suggest the possibility that hPDLSCs could also be used as an *in vitro* human cell-based model for neurogenesis and neuronal polarization [37]. In addition, the easy procedure for obtaining these from adults in normal or pathological condictions, may represent, as we have demonstrated with periodontal ligament cells from children [67,68], a suitable way of developing *in vitro* cell models of human diseases.

In this study, we also show that cell proliferation is not present through neurogenesis from hPDLSCs. The undifferentiated polygonal and fusiform cell shapes are reset and start their neuronal development as rounded spheres. The hPDLSCs-derived neurons gradually adopted a complex morphology by forming several processes, that grew and arborized, adquiring dendritic-like and axonal-like identities, giving rise to a variety of neuron-like morphologies. Futhermore, we may have discovered a transient cell nuclei lobulation coincident to *in vitro* neurogenesis, without being related to cell proliferation.

As noted above, morphological characteristics of the hPDLSCs, including cytoskeleton and nuclear morphology were examined in cells under proliferation and neural differentiation conditions. During interphase, the nuclei of proliferative hPDLSCs contained two or more nucleoli and the inside surface of the nuclear envelope is lined with the nuclear lamina. Furthermore, β-III tubulin microtubules are oriented parallel to the longitudinal axis of the cell. During mitosis, the nucleolus and the nuclear lamina are reversibly disassembled and β-III tubulin protein is present in the mitotic spindle. Moreover, mitotic chromosomes and cytokinesis were observed. Under neural differentiation conditions, the hPDLSCs displayed different nuclear shapes, including lobed nuclei connected by an internuclear bridge. Our results suggest the possibility that nuclear shape remodeling occurs during neurogenesis from hPDLSCs.

Morphological analysis revealed that cells with abnormal nuclei do not represent dividing cells due to the absence of cytokinesis, mitotic chromosomes and mitotic spindle during the described of *in vitro* neurogenesis processes or neuronal polarization from hPDLSCs. Futhermore, the nuclear lamina and nucleolus are not disassembled during *in vitro* neurogenesis from hPDLSCs, contrary to what happens during mitosis. Moreover, ultrastructural analysis with transmission electron microscopy revealed that hPDLSCs with abnormal nuclei do not represent apoptotic cells due to the absence of features of cells undergoing apoptotis (62).

Morphological analysis also revealed that the adult rodent V-SVZ of the anterolateral ventricle wall, as well as the SGZ of the hippocampal dentate gyrus, where adult neurogenesis has been clearly demonstrated, contains cells with nuclear shapes highly similar to those observed during *in vitro* neurogenesis from hPDLSCs.

Previous ultrastructure and immunocytochemistry studies also show that the V-SVZ stem cell niche contains cells with different morphologies and irregular nuclei [2–4,15–20]. Type-B cells have irregular nuclei that frecuently contain invaginations and irregular contours of the plasma membrane. Type-C cells nuclei contained deep invaginations and these cells are more spherical. Type-A cells have elongated cell body with one or two processes and the nuclei are occasionally invaginated [2]. Importantly, some studies have shown that murine and human V-SVZ have segmented nuclei connected by an internuclear bridge [18–20]. Although it has been suggested that lobed nuclei connected by an internuclear bridge are associated with quiescence in aNSCs [20], we observed that this kind of nuclei may be associated to nuclear movement within the cell during initial phases of neurogenesis, without being related to cell proliferation.

In addition, previous reports also shown irregular shaped nuclei in the adult SGZ [21–28]. Adult SGZ NSCs (type 1 cells) have irregular contours of the plasma membrane, and differences in heterochromatin aggregation has been also observed [9]. Futhermore, adult SGZ NSCs (type 2 cells) had an irregularly shaped nucleus [11,13]. Importantly, one study also found that many cultured hippocampal neurons have irregular nuclei or even consisted of two or more lobes connected by an internuclear bridge [29].

It has commonly been assumed that adult neurogenesis occurs progressively through sequential phases of proliferation [10,11]. Despite the advantages for the detection of adult neurogenesis using exogenous nucleotide analog administration or endogenous cell cycle markers, in addition to retroviral transduction, cell stage and lineage commitment markers, recent findings indicate that some observations interpreted as cell division could be false-positive signals [69–72]. The main method used to labed new neurons has been the incorporation of the thymidine analogs into the genome of dividing cells during S-phase of the cell cycle, but nevertheless thymidine analogs such as tritiated thymidine, BrdU, CldU and IdU may also be incorporated during DNA turnover or DNA repair [69–72]. Infection with retrovirus is another method used to label new neurons however, retroviral vectors not specifically infect dividing cells [71] and also, it has even been observed that postmitotic pyramidal neurons can also be labeled by fused infected microglia. [73]. Although the expression of endogenous cell cycle proteins is also used to label new neurons, recent findings indicate that cell cycle proteins expression is not necessarily related to cell division [72]. Proliferating cell nuclear antigen is also invoved in DNA repair [74]. Positivity of the proliferation marker KI-67 in noncycling cells has also been observed [75].

These findings indicate that there is a lack of a reliable definitive method to label new neurons. In addition, it is important to note that almost none of these labeled new born neurons show mitotic chromosomes or mitotic spindle to really confirm that adult neurogenesis occurs progressively through sequential phases of proliferation [2–9]. Moreover, the self-renewal and multipotent properties demonstrated by NSC in vitro [12] have not been clearly demonstrated in vivo [10,13,14].

Taken together, these results suggest the possibility that the sequence of events from aNSCs to neuron may also occur without being related to cell proliferation. It would therefore be interesting to examine whether SVZ and SGZ intermediate progenitor cells represent different stages of neurogenesis without being related to cell proliferation.

Beyond the central nervous system, the presence of lobed nuclei has been reported in most blood and immune cells, but the functional significance of multilobed nuclear structures is not yet known [76–79]. We observed that the nuclei of hPDLSCs during initial phases of neurogenesis are highly similar to those reported in immune cells. Thus, we suggest the possibility that multilobed nuclear structures may be associated to nuclear movement within the cell. Therefore, it would also be interesting to examine whether these putative madure cells also represent different stages of haematopoietic stem cell differentiation without being related to cell proliferation. Therefore, hPDLSCs could be also used to understand the functional significance of multilobed nuclear structures in blood and immune cells.

One of the most important discoveries in this work is the observation that small DNA containing structures may move within the cell to specific directions and temporarily form lobed nuclei. These small DNA containing structures displayed a spherical or ovoid shape, and it seems that some of them are connected to the main body of the nucleus by thin strands of nuclear material. It is important to note that some DNA containing structures are highly similar to nuclear envelope-limited chromatin sheets (ELCS) [20,80, 81]. Fibrillarin and laminin A/C proteins were detected in these small DNA containing structures.

It is known for many decades that chromatin particles can appear in the cellular cytoplasm and they are referred to as micronuclei, nucleoplasmic bridge and nuclear bud [82]. Although these nuclear anomalies have been associated with chromosomal instability events during mitosis [82–85], recent reports showed generation of micronuclei during interphase [86–88]. These findings call into question that micronuclei, nucleoplasmic bridge and nuclear bud does necessarily generated during mitosis [89]. Moreover, a high frequency of human mesenquimal stem cells with nuclear bud, micronuclei and nucleoplasmic bridge was detected under normal *in vitro* culture [90]. Therefore, the mechanisms that lead to extra-nuclear bodies formation and their biological relevance are still far from been understood [89,91,92].

In this study, we show that there can be a relationship in the formation of the nuclear bud, micronuclei, nucleoplasmic bridge and nuclear envelope-limited chromatin sheets. Taken together, these results suggest the possibility that the interphase cell nucleus may can reversibly disassembled into functional subunits that may moved independently within the cell, if necessary.

Multinuclear cells are commonly found in various human organs including heart, liver, salivary glands, muscle and endometrium, but their functional advantage remains unknown [93,94]. In addition, alterations in nuclear morphologies are closely associated with a wide range of human diseases, including muscular dystrophy and cancer [95,96].

hPDLSCs could facilitate an understanding of the mechanisms regulating nuclear morphology in response to cell shape changes and their functional relevance [78, 97].

## 5. Conclusion

hPDLSCs-derived neurons display a sequence of morphologic development highly similar to those reported before in primary neuronal cultures derived from rodent brains during neurogenesis, providing additional evidence that it is possible to differentiate hPDLSCs to neuron.

Cell proliferation is not present through neurogenesis from hPDLSCs, suggesting the possibility that the sequence of events from stem cell to neuron does not necessarily requires cell division from stem cell.

We may have discovered micronuclei movement and transient cell nuclei lobulation coincident to in vitro neurogenesis from hPDLSCs, suggesting the possibility that multilobed nuclear structures, commonly founded in various human organs, are associated to nuclear movement within the cell.

## Author Contributions

C.B. conceived of the study, designed the study, carried out the molecular lab work and drafted the manuscript. M.M. carried out the molecular lab work and participated in data analysis. S.M. conceived of the study, helped draft the manuscript and financial support. All authors discussed and commented on the manuscript

## Acknowledgments

We greatly appreciate the technical assistance of Microscopy Section of the University of Murcia in preparing samples for scanning electron microscopy and transmission electron microscopy.

---

**Figure S1.**
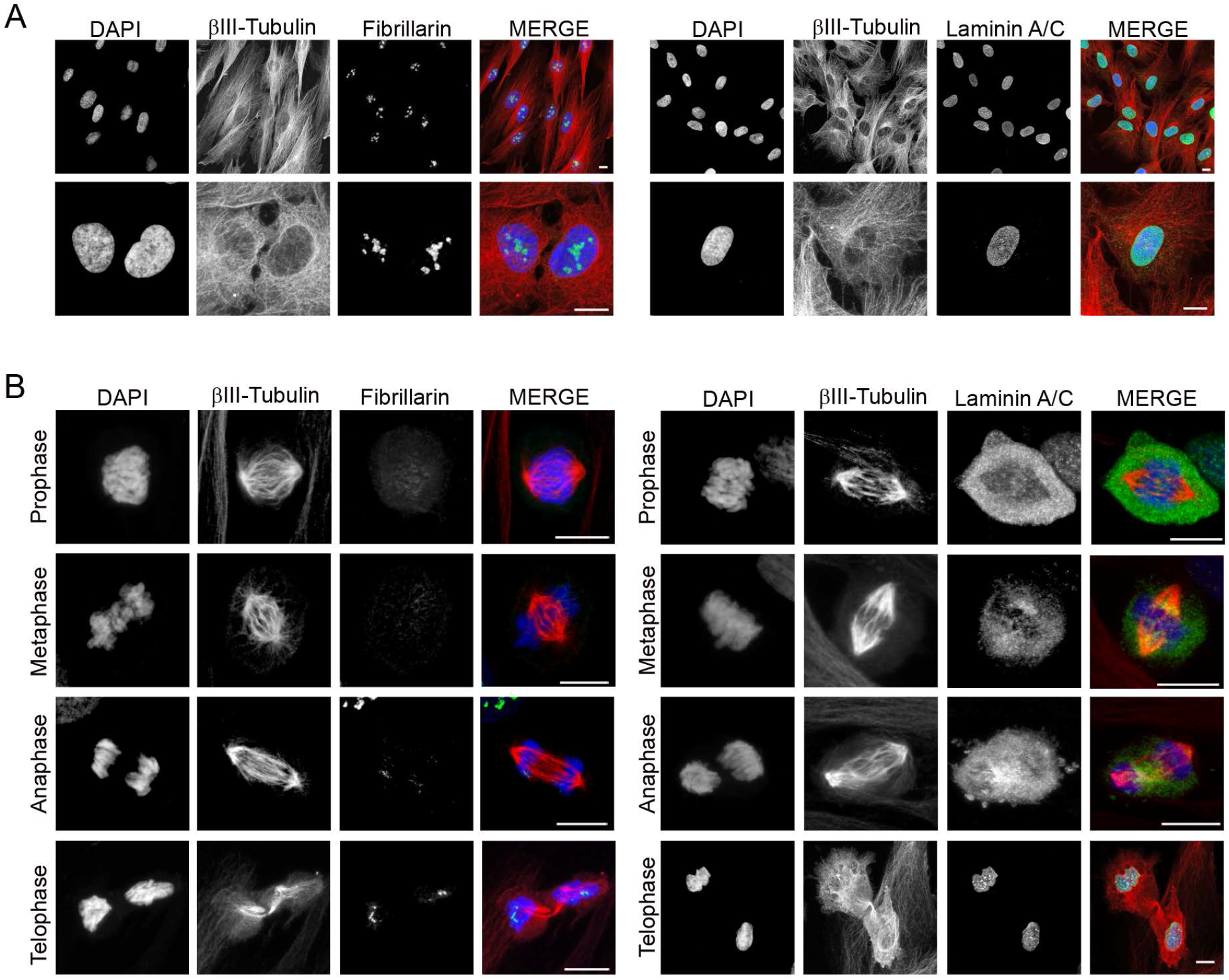
Dynamic localization of fibrillarin and laminin A/C proteins during the cell cycle of proliferative hPDLSCs. (**A**) During interphase, the nuclei of hPDLSCs contained two or more nucleoli and the inside surface of the nuclear envelope is lined with the nuclear lamina. (**B**) The nuclear lamina and nucleolus are reversibly disassembled during mitosis. Scale bar: 10 μm.

**Figure S2.**
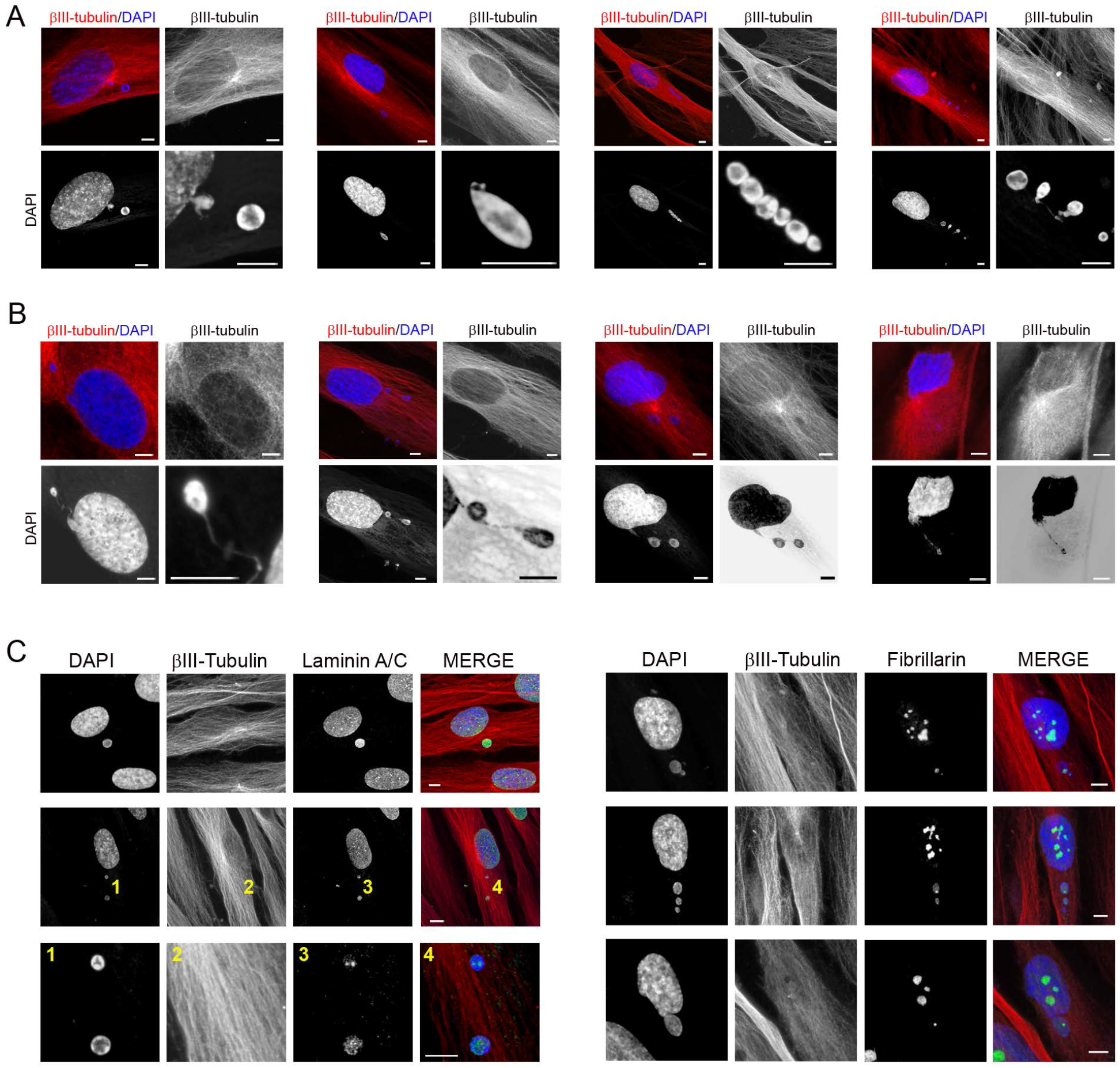
Cytoplasmic DNA containing structures. (**A**) Cytoplasmic DNA containing structures (micronuclei) displayed a spherical or ovoid shape and it seems that some of them are connected to the main body of the nucleus by thin strands of nuclear material (**B**). (**C**) Fibrillarin and laminin A/C proteins were detected in small DNA containing structures (numbers locate the areas showed in higher power). Scale bar: 5 μm.

**Figure S3.**
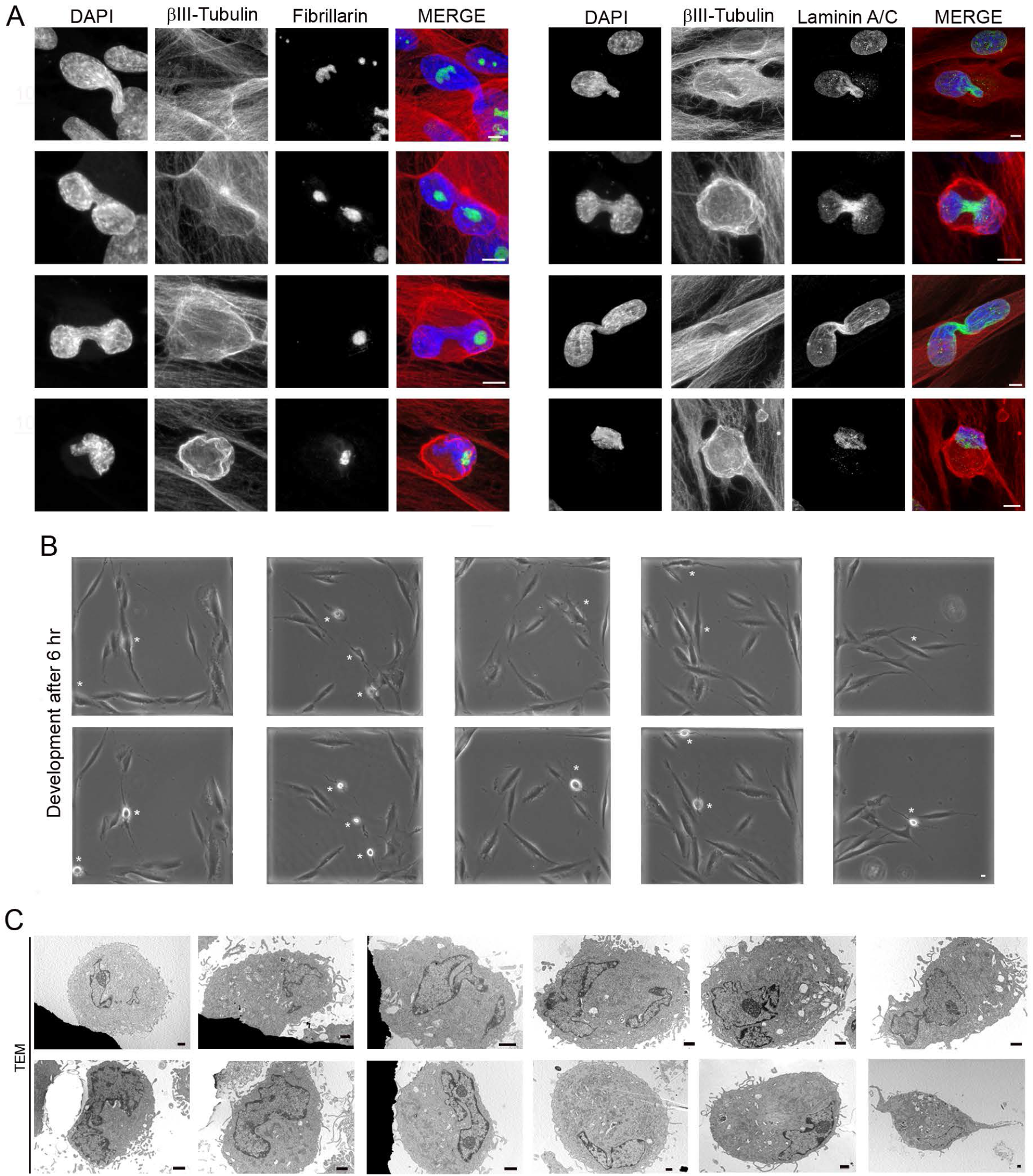
hPDLSCs-derived neurons are not directly generated through cell division from stem cells. (**A**) The nuclear lamina and nucleolus are not disassembled during in vitro neurogenesis from hPDLSCs. Futhermore, β-III tubulin is not present in the mitotic spindle. (**B**) Cytokinesis was not observed during in vitro neurogenesis processes (asterisks) from hPDLSCs. There is an interval of 6 hours between the micrographs. (**C**) Transmission electron micrographs show that hPDLSCs with abnormal nuclei do not represent apoptotic cells due to the absence of features of cells undergoing apoptosis. The scale bars are 5 μm in light microscope images and 1 μm in the transmission electron micrographs.

**Figure S4.**
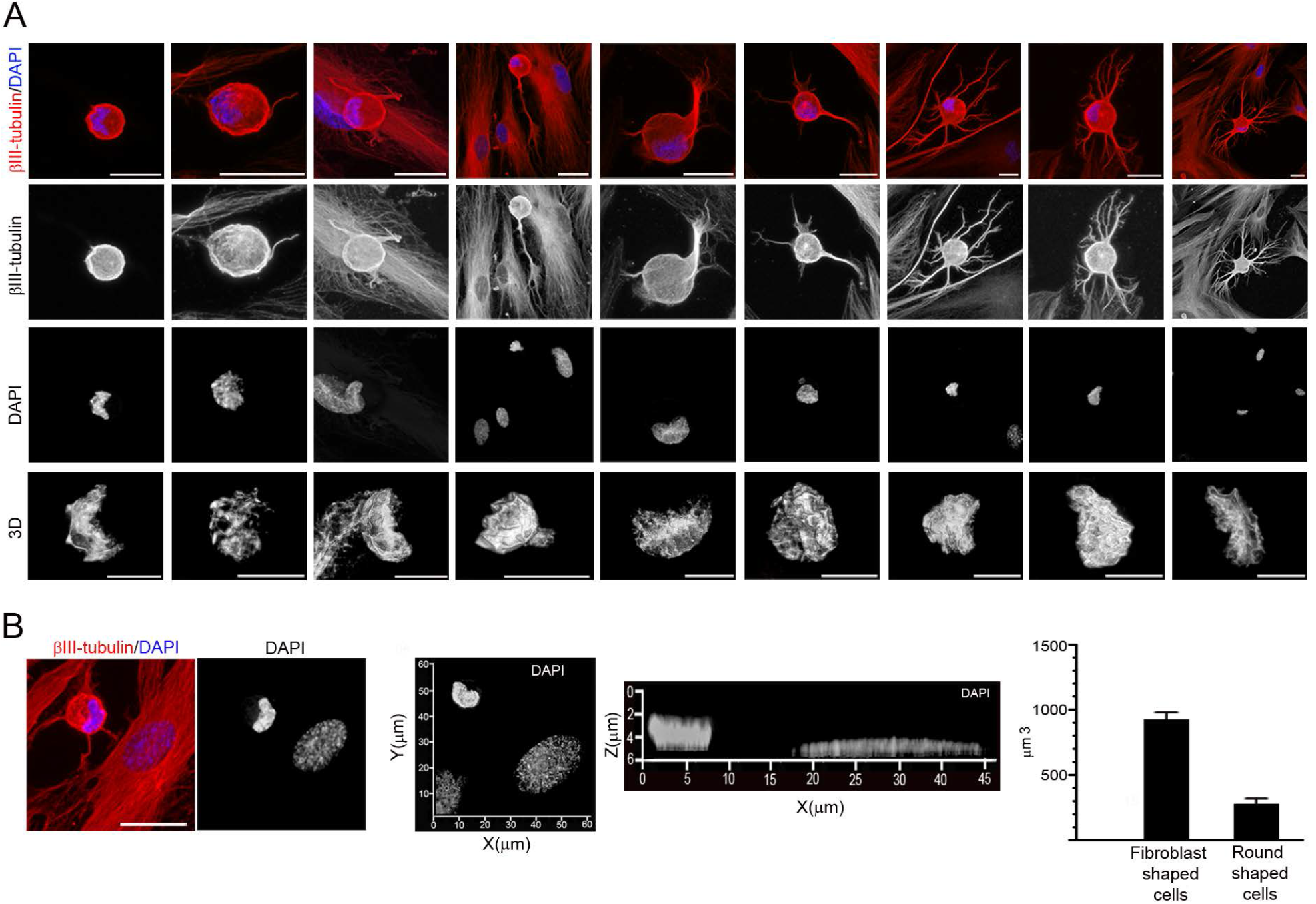
Nuclear shape in hPDLSCs-derived neurons during neuronal polarization. (**A**) No lobed nuclei are observed when hPDLSCs-derived neurons gradually acquired cellular polarity and more mature, neuronal-like morphology. (**B**) The nuclear volume shrinks as the cells become rounded during neurogenesis. Data represent mean ± S.E. of ten independent experiments. The scale bar in β-III tubulin and DAPI images are 50 μm and 10 μm for confocal 3D images of nuclei.

